# Socially-mediated compensatory growth carries hidden sperm costs in male guppies

**DOI:** 10.64898/2026.06.09.731059

**Authors:** Elisa Morbiato, Alexandra Glavaschi, Alessandro Devigili, Francesco Santi

**Affiliations:** Department of Biology, University of Padova, 35121 Padova, Italy; Institute for Vertebrate Biology, Czech Academy of Sciences, Brno, Czech Republic

## Abstract

Phenotypic plasticity allows organisms to mitigate early-life adversity through compensatory growth, yet the long-term costs of such “catch-up” trajectories remain poorly understood, particularly when driven by social factors. While most research focuses on nutritionally induced compensation, we investigate how early social competition—independent of resource availability—shapes adult life-history trade-offs in the guppy (*Poecilia reticulata*).

Our results show that scramble competition among peers during early development triggers compensatory growth once social constraints are removed. Crucially, we reveal a hidden reproductive cost: males exhibiting higher compensatory growth reach a similar adult size but produce significantly fewer sperm at maturity. This deficit persists even when controlling for adult body size, indicating a direct trade-off between somatic recovery and ejaculate investment. Furthermore, we find a negative association between gonopodium length and sperm count, suggesting competing allocations between pre- and postcopulatory traits during growth.

These findings reveal a cryptic cost of compensatory growth, where adult morphology conceals underlying differences in reproductive quality. By demonstrating that social environments alone can recalibrate life-history trajectories, we highlight the “ghosts of competition past” as critical determinants of fitness. Our study underscores the necessity of considering ontogenetic history to fully understand the evolution of sexually selected traits in social vertebrates.

## Introduction

Phenotypic plasticity, the ability of a single genotype to produce different phenotypes in response to environmental cues, is a fundamental mechanism shaping life-history strategies across the animal kingdom (1,2). An example of this plasticity is compensatory growth: a form of ontogenetic plasticity in which individuals of small body size in early life display elevated subsequent growth rates to reduce this initial size disadvantage (“catch-up growth”). This adaptive response has been widely documented across diverse taxa, including mammals, birds, and fishes, and is often modulated by factors such as resource availability, intra-group competition, and the social environment (3,4). While reaching the typical adult body size is beneficial for survival and fitness, theory predicts that such rapid, plastic growth responses carry physiological or reproductive costs because energy may be diverted away from somatic maintenance or reproduction (4,5).

One common ecological context in which such growth trajectories arise is early social competition (4). Early social competition occurs when developing individuals share an environment in which access to limited resources is constrained by the presence of conspecifics (6). Within this broader framework, peer competition represents a particularly important form of early social competition because peers repeatedly interact while exploiting the same pool of resources during development (7). Importantly, brood size, brood composition and asymmetries among siblings imply costs deriving from peer competition (8,9). In such cases, competition may approximate scramble competition, whereby individuals compete indirectly, through simultaneous exploitation of resources that cannot be monopolized, rather than through direct territorial exclusion or dominance interactions. Under scramble competition, small differences in body size or competitive ability can produce positive feedback between resource acquisition and growth, potentially leading to substantial divergence in developmental trajectories despite limited initial phenotypic variation (8).

Contrary to the assumption that peer competition is a mammalian-specific phenomenon, it is documented across diverse taxa, including birds (10), insects (11), and teleost fishes (12), where it serves to optimize individual fitness under resource limitation.

In the cost-benefits economy, trade-offs between growth and reproduction are expected to be higher in the sex experiencing stronger sexual selection, typically males (13). Intense male–male competition and female choice drive the evolution of costly traits that enhance reproductive success either before mating (precopulatory traits, such as ornaments and weapons) or after mating (postcopulatory traits, such as ejaculate characteristics) (14). In highly promiscuous species, where postcopulatory competition is intense, males are predicted to allocate proportionally more resources to ejaculate traits that enhance fertilization success. Conversely, when mating opportunities are limited, selection should favour greater investment in precopulatory traits that increase mating success (15).

Despite extensive work on both compensatory growth and reproductive trait plasticity, few studies have examined how early social environments influence growth trajectories and adult reproductive phenotype, especially in males (16–18).

The aim of this study is to explore how early social competition shapes growth rate, whether individuals exhibit compensatory growth once the stress is removed and if so, whether compensatory growth is associated with tradeoffs between pre and postmating traits.

The guppy (*Poecilia reticulata*) provides a powerful system to examine these allocation dynamics. Guppies are small freshwater fish native from Central America, with internal fertilisation and indeterminate growth (19). Male guppies exhibit conspicuous carotenoid-based orange coloration that enhances mating success and is strongly condition-dependent (20–22). Moreover, male guppies possess a gonopodium, a modified anal fin that functions as an intromittent organ for internal fertilization, and variation in gonopodium length has been shown to influence male mating and reproductive success (20,22). This species is characterized by high levels of polyandry and intense sperm competition (23), resulting in substantial allocation to ejaculate production.

Importantly, sexually selected traits in guppies are not only costly but also plastically expressed. Males adjust ejaculate characteristics in response to social cues: individuals exposed to females produce faster sperm than those deprived of female stimuli, indicating adaptive plasticity in sperm quality according to perceived mating opportunities (24). However, high investment in sperm production is associated with physiological costs, including reduced telomere length in germinal and somatic tissues (25), consistent with trade-offs between reproductive allocation and somatic maintenance.

In guppies, the social environment experienced during the juvenile phase can shape life-history, including the timing of development of male secondary sexual traits (12,26,27), and offspring size itself has been interpreted as an adaptation to the competitive environment encountered by newborn (12).

Together, these studies suggest that early social interactions may generate meaningful variation in resource acquisition during development, with downstream consequences for growth trajectories and later life-history allocation. However, the extent to which early social environments modulate compensatory growth, and whether such growth entails measurable reproductive costs, remains largely untested in social vertebrates.

To fill this gap, we reared guppies with their broods during early -life and subsequently in isolation, and quantified compensatory growth using both longitudinal and cross-sectional approaches. During the first 60 days after birth, fry were maintained within their natal broods, generating natural variation in competition arising from brood size differences. Although brood size was not experimentally manipulated, housing entire broods in confined environments created variation in competition for food and space, conditions known to influence growth trajectories in juvenile fishes (3,12,28). Competition for resources among juvenile guppies is likely to approximate a scramble competition scenario, where individuals exploit shared, limited resources simultaneously and success depends on rapid resource acquisition rather than dominance (7,8). Hence, natural variation in size among individuals may generate unequal access to food and space within broods, in turn biasing growth trajectories over time. The intra-brood competition should be interpreted within the broader context of peer competition occurring among individuals of similar age and size within population, where size-structured interactions among social species can generate similar competitive dynamics beyond the brood level. Indeed, evidence from fish species suggests that social competition among peers results in foraging hierarchies of local resources (29).

Our design therefore captures developmental variation mediated by the social environment without imposing artificial constraints beyond those experienced in natural brood contexts (30). This preserves ecologically relevant heterogeneity in the social environment and among-individual developmental trajectories, thereby improving the ecological realism of the experiment rather than obscuring it by artificially minimizing meaningful variation (31).

We individually isolated males at 60 days because sexual traits become visible at this age (32). Isolation removed direct intra-brood competition and enabled us to quantify growth responses conditional on prior developmental state. We measured sexual traits at 120 days, corresponding to full sexual maturity in males, when sexual traits are fully expressed and can be accurately assessed (33).

We predicted: (i) early differences in body size would predict divergent growth trajectories under social competition; (ii) males exhibiting reduced growth during competitive social rearing would subsequently display compensatory growth in isolation; (iii) compensatory growth would be associated with reduced investment in sexually selected traits.

## Materials & Methods

### Stock population

Guppies used in this experiment were derived from a semi-natural population established at the Botanical Garden of Padova in 2014. The founders originated from a stock collected in 2003 from the Lower Tacarigua River in Trinidad. All fish were acclimated at the University of Padova in 150-L tanks with an equal sex ratio and a density of 1 fish/L, under a 12 h light:dark photoperiod and water temperature of 24–27°C. Fry were collected monthly from stock tanks and separated into single-sex tanks at sexual maturation to obtain virgin females and males of known age.

### Experimental overview

We obtained experimental fish by pairing adult males (between 4 and 6 months of age) with virgin females and allowing them to mate for 3 days. Afterwards, females were individually isolated in 3.5-L tanks equipped with nurseries and checked twice daily until they produced a brood, or for a maximum of 60 days. In total, 93 broods were obtained from 93 mothers and 50 fathers. Each brood was uniquely associated with a single mother, whereas sires were represented across multiple broods: 21 fathers sired offspring with one mother, 19 with two mothers, 6 with three mothers, and 4 with four mothers.

We photographed broods within 72 h of birth and then maintained them in 3.5-L tanks until 60 days of age. Because individuals were reared together with their brood, they experienced continuous social competition for food and space; we therefore defined the period between day 3 and 60 as the “social competition treatment”. During this time, all fish were fed *ad libitum* with dry food and

### *Artemia* sp. nauplii twice daily

At 60 days of age, we randomly selected up to 3 males per brood, which were photographed and isolated in clear perforated cylinders (10 cm diameter) within 120-L tanks containing 30 cm of water, 2 cm of gravel, a mechanical filter, 10–15 free-swimming females (as stimuli to favour regular development of sexual traits), and an aquarium plant. Density was limited to a maximum of 20 males (i.e. cylinders) per tank. This setup allowed individual recognition while removing competition for food and space. During this period (henceforth “the social isolation treatment”), fish were checked and fed daily an *ad libitum* diet consisting of dry food and *Artemia* nauplii twice. The isolation lasted for 60 days, and at day 120 all males were photographed and stripped of sperm for the purpose of count and velocity measurement (see below).

Our design integrates both an early cross-sectional assessment of growth under sibling competition, and a later longitudinal phase in which individual growth and reproductive traits could be quantified after isolation. This approach allows us to separate developmental effects emerging under peer competition from later individual growth responses once social constraints are removed.

### Body size and precopulatory traits measurement

We photographed broods within 72 h of birth in a Petri dish on grid-lined paper under a stereomicroscope (Leica S59C, Leica Microsystems, Wezlar, Germany). We then estimated individual total length from the photos using ImageJ. During this early developmental phase, as individuals were maintained together with their brood, it was impossible to keep track of individual identity. To account for the lack of individual identification at day 3, we assigned each male’s initial size by randomly sampling from the pool of total length measurements taken from its own brood at day 3. We validated this stochastic assignment by calculating both the Intraclass Correlation Coefficient (ICC) and the mean within-brood Coefficient of Variation (CV) for total length at day 3. We observed high phenotypic homogeneity within broods at day 3 (Adjusted ICC = 0.74; mean within-brood CV = 3.04). The high ICC shows that 74.4 % of body size variation at day 3 is attributable to brood membership, while the low within-brood CV (3.04 %) shows that individual sizes deviate minimally from brood averages. Together, these metrics demonstrate that random size assignment at day 3 provides a robust proxy for individual initial size, allowing us to both preserve the observed intra-brood variability while remaining consistent with the high resemblance among siblings.

At day 60, actual individual identities became available. At both 60 and 120 days of age males were lightly anesthetized using MS-222 and photographed on their left side on a grid-lined paper under a stereomicroscope (Leica S59C, Leica Microsystems, Wezlar, Germany). From these photos, we measured standard length, gonopodium length, and the area of orange spots relative to body area using ImageJ.

### Relative growth rate

To quantify growth trajectories, we calculated the relative growth rate (RGR) between day 3 and day 60 (RGR_3_–_60_) and between day 60 and day 120 (RGR_60_–_120_) as:

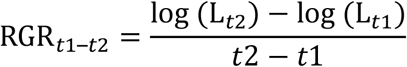

where L is the body length (mm) measure recorded at each time point. Specifically, L corresponded to total length at day 3, and to standard length at days 60 and 120, when males were measured individually. At day 3, total length was used instead of standard length because the caudal fin is still poorly differentiated at this stage. RGR was calculated using log-transformed length, reflecting the multiplicative nature of somatic growth and allowing growth to be expressed on a proportional scale. Log-transformation standardizes growth rates with respect to initial size, such that individuals showing equivalent proportional increases in length have comparable RGR values regardless of their starting size (Dmitriew 2011). This metric captures size-standardized growth during the social and isolation phases and allows direct comparison among individuals that differed in early size.

### Ejaculate assays

We measured sperm production and sperm velocity for all males at day 120 using standardised procedures for *Poecilia reticulata* (see Supplementary Methods for details). Sperm count was estimated from the number of sperm bundles while sperm velocity was quantified using computer-assisted sperm analysis (CASA) (33).

## Statistical analyses

### Cross and longitudinal growth analysis

Covariation between growth rates was assessed by setting RGR_60_–_120_ as the dependent variable as a function of RGR_3_–_60_. Maternal and paternal identities were included as random intercepts. Models were fitted using restricted maximum likelihood estimation (REML), and t-tests were computed using Satterthwaite’s method.

To characterize growth trajectories across development, we calculated RGR separately for the two treatments and fitted linear mixed-effects models. RGR was set as the dependent variable, and log-transformed total length at 3 days, brood size, and their interaction as predictors. Initial size was included to account for size-dependent growth, while brood size was included to capture potential effects of the social environment on growth trajectories. To account for shared genetic and environmental effects, maternal and paternal identities were included as random intercepts. Models were fitted using restricted maximum likelihood estimation (REML), and t-tests were computed using Satterthwaite’s method.

### Relationships between growth rate and sexual traits

We assessed the relationship between RGR_60_–_120_ and sperm number and velocity (Average Path Velocity: VAP), gonopodium length, and orange coloration, to identify potential associations of investment in sexual traits using linear mixed-effects models.

Since each offspring was associated to both its mother and its father, we included both maternal and paternal identity as random intercepts to account for shared genetic and environmental effects. Brood size, VAP, orange coloration and gonopodium length were z-score transformed (mean = 0, SD = 1) to facilitate comparison of effect sizes. Estimates are reported with standard errors and Satterthwaite-approximated p-values. Model assumptions were evaluated using residual diagnostics.

We further investigated the relationship between social isolation, growth rate, and reproductive investment by conducting a separate ordinary least squares (OLS) regression. Log-transformed sperm number was modelled as a function of RGR_60_–_120_, while controlling for adult body size (log-transformed SL at 120 days). Gonopodium length at 120 days was included as an additional covariate. This OLS approach provided a direct estimate of the effect of compensatory growth on sperm production and gonopodium length. Model assumptions were evaluated using residual diagnostics.

Sample sizes differed between analyses due to missing measurements at specific developmental stages or natural mortality. Relative growth rate during the social competition phase (RGR_3_–_60_) was calculated for 89 individuals with complete body size measurements at both day 3 and day 60. For the social isolation phase (RGR_60_–_120_), we fitted two complementary linear mixed-effects models addressing different biological questions. First, to test the overall effect of the early social environment on subsequent catch-up growth, we fitted a simplified model including brood size as a predictor, which retained 109 individuals with measurements at both day 60 and day 120. Second, to test whether compensatory growth depended on developmental state at birth, we fitted an interaction model including body size at day 3, brood size, and their interaction. Because this analysis required complete longitudinal measurements across all three time points (days 3, 60, and 120), the sample size was restricted to 66 individuals. Linear mixed-effects models examining associations between RGR_60_–_120_ and male sexual traits included individuals with complete data for RGR and all focal traits (sperm number, sperm velocity, gonopodium length, and orange coloration; n = 98). Ordinary least squares regression of sperm number included all individuals with available RGR_60_–_120_, standard length, and gonopodium length measurements (n = 98). This approach allowed each analysis to retain all individuals with complete data for the variables of interest while accounting for missing measurements and mortality.

All analyses were conducted in R (version 4.4.2), using the packages *lme4, lmerTest, performance, emmeans*.

## Results

### Early growth is negatively associated to later growth

RGR_60_–_120_was negatively associated with RGR_3_–_60_, indicating that variation in growth during the social competition period was inversely related to growth during social isolation, consistent with compensatory growth hypothesis. In a mixed-effects model including Father ID and Mother ID as random intercepts RGR_3_–_60_ significantly predicted RGR_60_–_120_ (β = –0.329, p < 0.001).

### Early body size and brood size shape growth trajectories

Male guppies exhibited substantial inter-individual variation in RGR during both social (3–60 days; RGR range: 0.00685 – 0.0176, SD = 0.00187) and isolation (60–120 days; RGR range: -0.00090 – 0.00723, SD = 0.00145) developmental periods. Mixed-effects models revealed that total length at birth and brood size influenced RGR, with maternal identity contributing additional variance.

Between 3 and 60 days, total length had a significant negative effect on RGR (p < 0.001), indicating that individuals of larger size at birth grew more slowly during the social competition phase, consistent with accelerated growth in smaller individuals. Random effects were extremely small, indicating that differences among families contributed very little to growth variation at this stage (Table 2, Figure 2).

**Table 1.** RGR_3_–_60_ negatively predicts RGR_60_–_120_. Linear mixed-effects model testing whether RGR_3_–_60_ predicts RGR_60_–_120_. Mother ID and Father ID were included as random intercepts. Estimates are reported with standard errors, Satterthwaite-approximated degrees of freedom, t-values, and p-values. Variance components for random effects are shown.

| Fixed effects | Estimate | s.e. | d.f. | t | p |
| --- | --- | --- | --- | --- | --- |
| Intercept | 0.0064 | 0.0011 | 65.21 | 5.862 | <0.001 |
| $RGR_{3-60}$ | -0.3289 | 0.0919 | 65.76 | -3.578 | <0.001 |
| Random effects | Variance | s.d. |  |  |  |
| Mother ID | $7.15 \times 10^{-7}$ | 0.00085 | | | |
| Father ID | $3.54 \times 10^{-7}$ | 0.00060 | | | |

**Table 2.** Body size at birth negatively predicts RGR_3_–_60_. Linear mixed-effects model testing the effects of body total length at 3 days, brood size, and their interaction on early RGR (RGR_3_–_60_). Mother ID and Father ID were included as random intercepts. Estimates are reported with standard errors, Satterthwaite-approximated degrees of freedom, t-values, and p-values. Variance components for random effects are shown.

| Fixed effects | Estimate | s.e. | d.f. | t | p |
| --- | --- | --- | --- | --- | --- |
| Intercept | 0.0446 | 0.0036 | 69.58 | 12.532 | <0.001 |
| TL<sub>3</sub> | -0.0173 | 0.0019 | 69.42 | -9.235 | <0.001 |
| Brood size (z) | -0.0039 | 0.0034 | 78.98 | -1.178 | 0.242 |
| TL<sub>3</sub> × Brood size (z) | 0.0017 | 0.0018 | 79.71 | 0.941 | 0.350 |
| Random effects | Variance | s.d. |  |  |  |
| Mother ID | $7.59 \times 10^{-7}$ | 0.00087 | | | |
| Father ID | $6.03 \times 10^{-7}$ | 0.00078 | | | |

**Figure 1.**
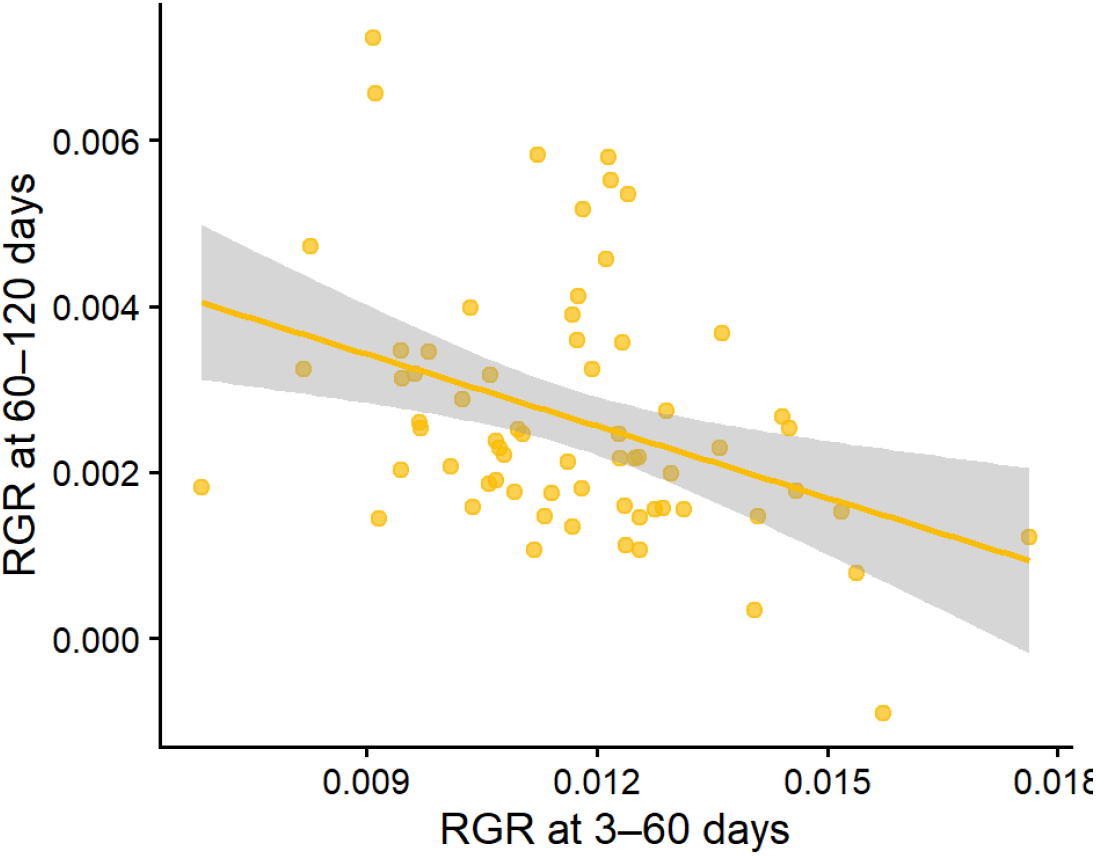
RGR_3_–_60_ negatively predicts RGR_60_–_120_. Relationship between RGR_3_–_60_and RGR_60_–_120_. Each point represents an individual. The solid yellow line shows the fitted linear regression with 95% confidence intervals (grey). The negative slope indicates that individuals with lower social growth rates exhibited higher growth during the isolation phase, consistent with compensatory growth.

**Figure 2.**
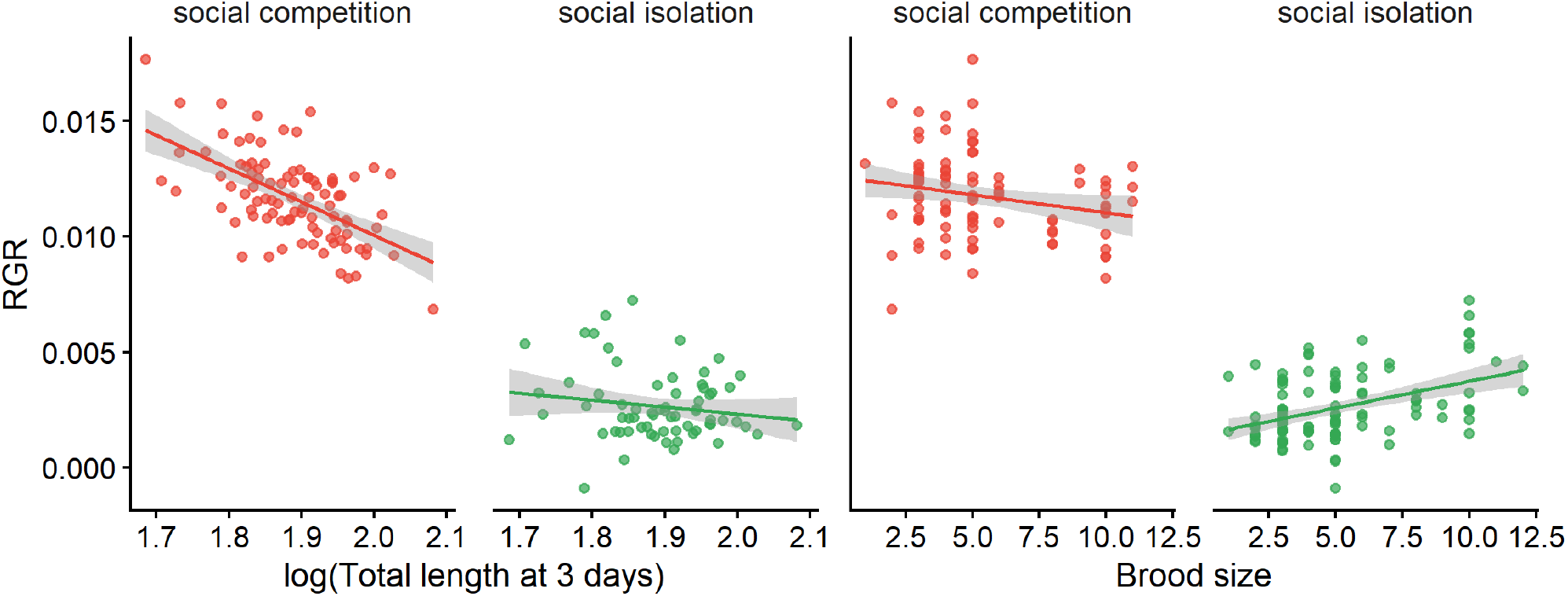
Accelerated growth depends on social environment. Left panel: relationship between total length at 3 days and Relative Growth Rate (RGR). During the social competition phase (3–60 days, in red), offspring that were smaller exhibited higher growth rates, consistent with accelerated growth. During the subsequent isolation phase (60–120 days, in green), this relationship disappeared, indicating that accelerated growth was curtailed once social competition was removed. Right panel: Relationship between brood size and relative growth rate (RGR) during the social competition phase (3-60 days; red) and the social isolation phase (60-120 days; green). Brood size did not predict RGR during the social phase, but was positively associated with RGR after isolation. Lines show fitted linear regressions with 95% confidence intervals.

Between 60 and 120 days, brood size had a positive significant effect on RGR (p = 0.002), and the interaction between total length at 3 days and brood size was negative and significant (p = 0.004), indicating that individuals from larger broods who were larger at birth grew slower during the social isolation phase. Both maternal and paternal identity contributed negligible variance, highlighting that the majority of variability in growth during isolation was explained by individual-specific factors rather than family-level effects. (Table 3, Figure 2).

**Table 3.** Body size and brood size interaction negatively predicts RGR (RGR_60_–_120_). Linear mixed-effects model testing the effects of body total length at 3 days, brood size, and their interaction on post-social RGR (RGR_60_–_120_). Mother ID and Father ID were included as random intercepts. Estimates are reported with standard errors, Satterthwaite-approximated degrees of freedom, t-values, and p-values. Variance components for random effects are shown.

| Fixed effects | Estimate | s.e. | d.f. | t | p |
| --- | --- | --- | --- | --- | --- |
| Intercept | 0.0008 | 0.0039 | 47.07 | 0.194 | 0.847 |
| TL<sub>3</sub> | 0.0008 | 0.0021 | 47.21 | 0.400 | 0.691 |
| Brood size (z) | 0.0105 | 0.0036 | 51.80 | 2.939 | 0.004 |
| TL<sub>3</sub> × Brood size (z) | -0.0051 | 0.0019 | 51.69 | -2.716 | 0.008 |
| **Random effects** |  |  |  |  |  |
|  | Variance | s.d. |  |  |  |
| Mother ID | 3.08 × 10<sup>-7</sup> | 0.00056 |  |  |  |
| Father ID | 0 | 0 |  |  |  |

### Compensatory growth predicts reduced sperm production

Among all the pre- and postcopulatory sexual traits considered in our analyses, only sperm production had a significant association with relative growth rate (Tables S1-3). Indeed, sperm production at 120 days was strongly associated with prior growth history. In the OLS model, RGR_60_–_120_ had a significant negative effect on sperm number (p < 0.001), indicating that males exhibiting higher growth during this period produced fewer sperm at sexual maturity (Table 4, Figure 3).

**Table 4.**
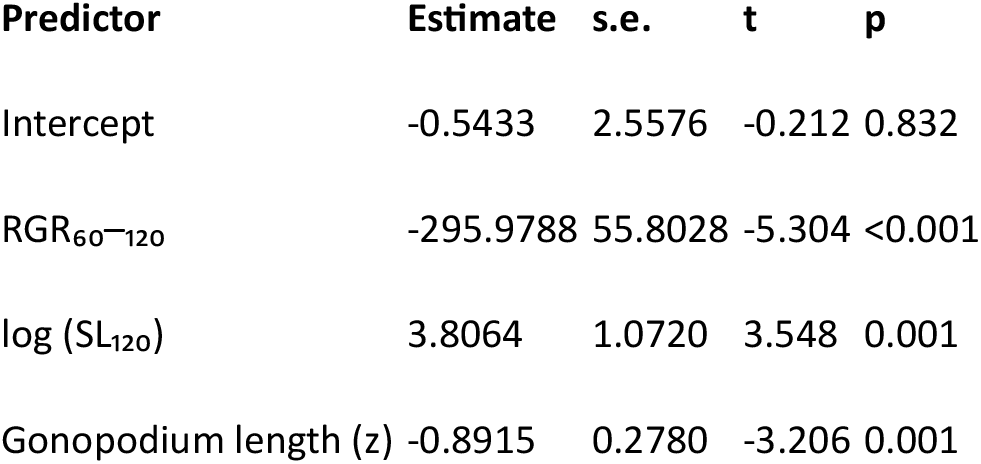
Compensatory growth negatively predicts sperm number. Ordinary least squares regression testing whether social isolation RGR (RGR_60_–_120_) predicts male reproductive investment, measured as log-transformed sperm number. Standard length and gonopodium length at 120 days were included as covariates. Regression coefficients with standard errors, t-values, and p-values are reported.

| Predictor | Estimate | s.e. | t | p |
| --- | --- | --- | --- | --- |
| Intercept | -0.5433 | 2.5576 | -0.212 | 0.832 |
| $RGR_{60-120}$ | -295.9788 | 55.8028 | -5.304 | <0.001 |
| $\log(SL_{120})$ | 3.8064 | 1.0720 | 3.548 | 0.001 |
| Gonopodium length (z) | -0.8915 | 0.2780 | -3.206 | 0.001 |

**Figure 3.**
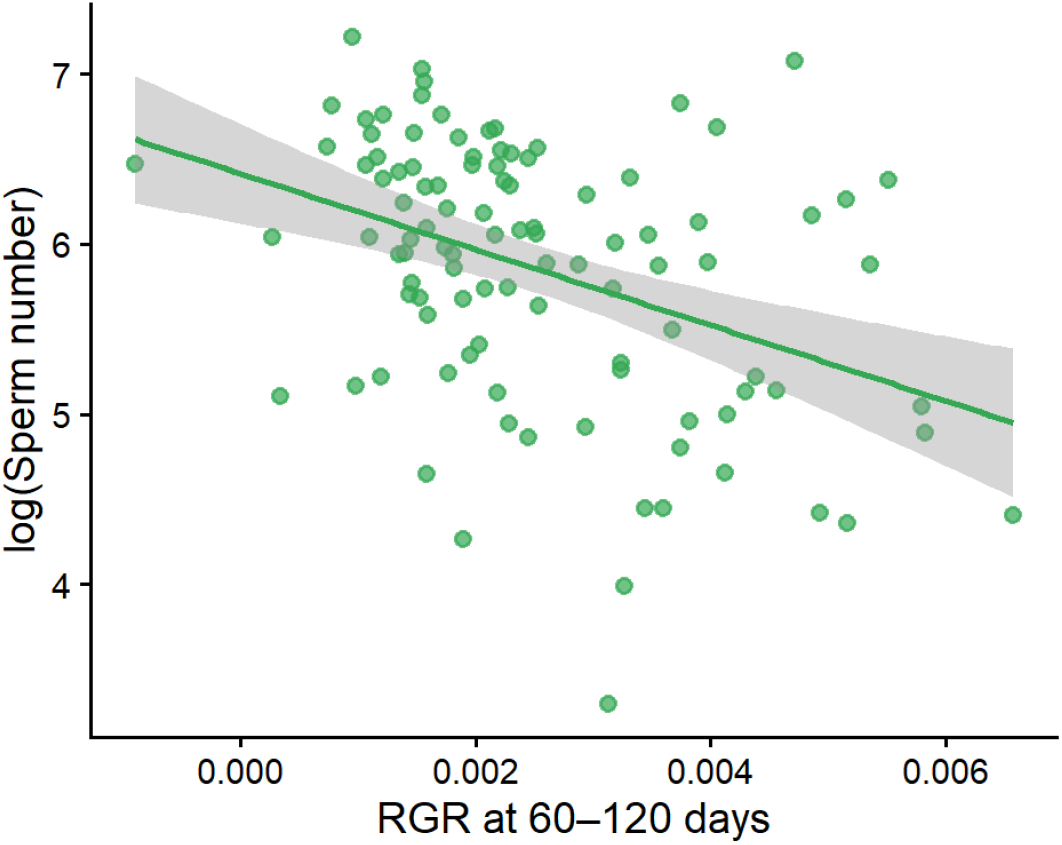
Compensatory growth negatively predicts sperm production. Relationship between RGR_60_–_120_ and sperm number. Males that exhibited higher RGR_60_–_120_ produced fewer sperm as adults. The regression line represents the effect of RGR estimated from an OLS model controlling for adult body size (standard length at 120 days) and gonopodium length.

Standard length at 120 days of age positively predicted sperm production (p = 0.001) while gonopodium length showed a significant negative association with sperm number (p = 0.001), suggesting a trade-off between investment in pre and postcopulatory sexual traits. The model explained a substantial proportion of the variance in sperm number (adjusted R^2^ = 0.27).

## Discussion

### Social competition as a driver of growth plasticity

We found that body size at day 3 was strongly negatively associated with the growth rate between 3 and 60 days, whereas later (60–120 days) this relationship disappeared. This suggests that growth plasticity is socially mediated: within-group competition promotes accelerated growth in smaller individuals, whereas isolation attenuates it.

Variation in standard length observed at the onset of social isolation (60 days) provides a measure of individual developmental state shaped by social competition, which subsequently predicts individual growth responses and reproductive investment in adulthood.

While previous studies often induced growth compensation by delaying sexual maturation through caloric restriction (34,35), our results show that social competition—as the scramble competition for space and resources—is sufficient to trigger these plastic trajectories. This aligns with the “social competition hypothesis,” where the mere presence of peers acts as a primary driver for phenotypic and life-history variation, independent of resource availability (36).

### Trade-offs between compensatory growth and reproductive allocation

The most striking finding is the cost of growth compensation on sperm production. Even though males with high RGR_60_–_120_ partially or fully compensated for their initial size disadvantage, they paid off compensatory growth with significantly lower sperm reserves. This supports the theory that rapid growth is energetically demanding and diverts resources away from reproduction or maintenance (5). The trade-off between somatic recovery and ejaculate investment remains significant when controlling for adult body size suggesting that it is not simply a matter of “small fish produce less sperm” (37) but rather a specific cost of the rate of growth itself (25).

This finding is supported by a growing body of evidence showing that across taxa ejaculate traits are highly sensitive to nutritional and developmental history (38). In mosquitofish, a restricted early diet leads to lower sperm reserves and replenishment rates, although these effects may attenuate with age (34). Such trade-offs between rapid juvenile growth and adult reproductive investment have been documented in other poeciliids (39,40) and are thought to reflect life-history constraints where early growth must be balanced against future reproductive output. Our study shows that even when resources are not limited, the costs of scramble competition with peers persist over time, and the metabolic demands of compensatory growth are still able to create a similar deficit in sperm production.

Unlike precopulatory traits (like orange coloration, which showed no significant association with RGR in our study), ejaculate production is a continuous, resource-intensive process in guppies. As such, its cost is predicted to be higher than other sexual traits (25).

We also observed a significant negative association between gonopodium length and sperm number. This suggests a trade-off between investment in precopulatory sexual traits and ejaculate production (37). While fighting for dominance or territory has been shown to have negligible life-history costs in some poecilids (41), the direct cost of sperm production remains high. Males that prioritize somatic growth and the development of morphological traits (like the gonopodium) may be severely disadvantaged in sperm competition scenarios, where higher sperm numbers are the primary determinant of paternity success (33).

### Evolutionary implications for postcopulatory success

In highly promiscuous mating systems characterised by intense sperm competition, a reduction in sperm number likely translates to a direct loss of fitness. Our results suggest that males that catch up in size may be trading off future competitive fertilization success for the immediate benefits of being larger such as better survival or enhanced precopulatory attractiveness. This highlights how early-life social environments can create cryptic cost in compensatory growth where an individual appears robust in size but is functionally weaker in reproductive quality. Hence, two males of identical adult size may have vastly different reproductive potential based on their ontogenetic path. This “hidden” cost of early social competition—reduced sperm production—could potentially reduce paternity share and lifetime fitness, even if the male appears phenotypically “recovered” in adulthood. This underscores the importance of considering early social environments as critical determinants of adult reproductive allocation. In fact, in many evolutionary studies, adult body size is treated as a reliable proxy for individual quality or resource-holding potential (42). However, our results demonstrate that this static view of the phenotype can be profoundly misleading due to the phenotypic masking of compensatory growth (43). Consequently, studies focusing solely on adult body size as proxy of fitness may fail to capture the hidden costs of early-life social interactions.

We show that intra-brood competition acts as a powerful developmental filter. In nature, peer competition is a ubiquitous form of scramble competition where access to resources is dictated by subtle asymmetries in birth size and social density (8). Our data indicate that the stress of early social interactions is sufficient to trigger long-term trade-offs, even when food is subsequently provided *ad libitum*.

Similar long-term trade-offs from mammals indicates that early-life sibling competition can generate persistent effects on adult reproductive phenotype. In deer mice, males reared with more brothers develop larger testes and increased reproductive output, consistent with early-life tuning of sperm production in response to perceived competitive environment (44). Similarly, experimental work in house mice shows that early social conditions can induce plastic changes in sperm production and testicular function (45). More broadly, litter composition and sibling competition are known to structure developmental trajectories in mammals, often producing long-term effects on physiology and life-history traits (46).

In conclusion, we demonstrate that socially-mediated compensatory growth can mask underlying differences in quality. In highly promiscuous mating system, where sperm number is a key determinant of fertilization success, social competition could determine lower paternity share in competitive scenarios by reducing the allocation in sperm production to favour growth. This underscores the importance of considering ontogenetic history as a critical determinant of adult reproductive allocation in social vertebrates.

## Ethics

The ethic protocol applied to this study was approved by the University of Padova Institutional Ethical Committee (permit no. 256 /2018).

## Data accessibility

Scripts and datasets are publicly available in Zenodo: https://doi.org/10.5281/zenodo.20597163.

## Declaration of AI use

The authors acknowledge the use of AI-assisted technologies for supporting text drafting and linguistic refinement. These tools were used under strict human oversight; all content was independently verified, critically assessed, and modified where necessary prior to submission. Responsibility for the integrity of the work rests solely with the authors.

## Author’s contributions

This paper is part of a larger study on male guppy reproduction and longevity which was conceived by Andrea Pilastro. E.M. conceived the present study. E.M. and A.G. conducted the experimental work. A.D. and F.S. contributed to data collection, and F.S. performed the morphological analyses. E.M. carried out the statistical analyses. E.M. wrote the manuscript with contributions from all authors. All authors approved the final version of the manuscript.

## Acknowledgements

We thank Andrea Pilastro for providing laboratory facilities and resources. We would also like to thank Milan VrtÍlek for his insightful comments and valuable discussions on an earlier version of this manuscript. We are grateful to the staff of the University of Padua fish facility for their expert assistance with animal husbandry.

## Supplementary Information

### Ejaculate assays

At 120 days of age, males were anesthetized in an MS-222 bath and stripped of sperm immediately after being photographed (see above). Fish were placed under a dissection microscope, the gonopodium swung back and forth and sperm released in a drop of 1% saline solution by applying gentle pressure to the abdomen. Guppy ejaculates are organized in discrete packages (bundles), each containing roughly 21000 sperm cells (Boschetto et al 2011). Ejaculates were photographed with a Canon 450D camera mounted to a light microscope and sperm bundles were counted from pictures using ImageJ. Sperm velocity (µm/s) was measured with a CASA system (Hamilton-Thorne). Three sperm bundles were placed on a multiwell slide coated in 1% polyvinyl acid to prevent adherence, and activated in a medium containing 150 mM KCl and 2 mg/L bovine serum albumin. Velocity was measured as cells swam away from the opening bundle. The sperm tracker produces three measures of velocity (curvilinear, straight line and average path) which are closely corelated in guppies (Gasparini et al 2010). We retained average path velocity (VAP) for analyses.

### RGR relationship with pre and postcopulatory traits

We used multivariate linear mixed-effects models to examine whether RGR_60_–_120_ was associated with male reproductive traits measured at 120 days of age. Among the examined traits, only sperm number showed a statistically significant association with RGR_60_–_120_, whereas sperm velocity, coloration, and gonopodium length showed non-significant effects even after removing the least significant predictors in reduced models (Table S1, Table S2, Table S3).

**Table S1.**
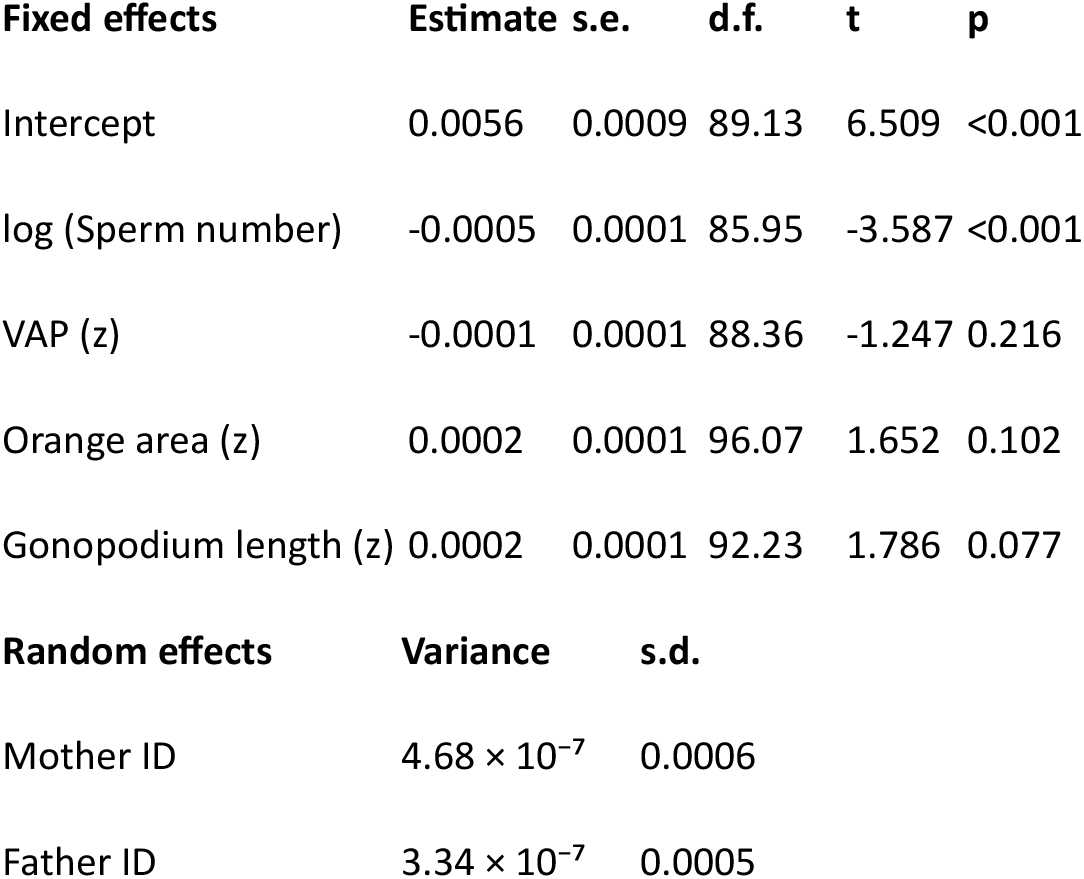
Linear mixed-effects model testing the association between social isolation RGR (RGR_60_–_120_) and male sexual traits measured at 120 days of age. Fixed effects include log-transformed sperm number, sperm velocity (VAP), orange coloration area, and gonopodium length. Maternal identity (Mother ID) and paternal identity (Father ID) were included as random intercepts. All continuous predictors were standardized (mean = 0, SD = 1). Estimates are reported with standard errors and Satterthwaite-approximated *p*-values.

**Table S2.**
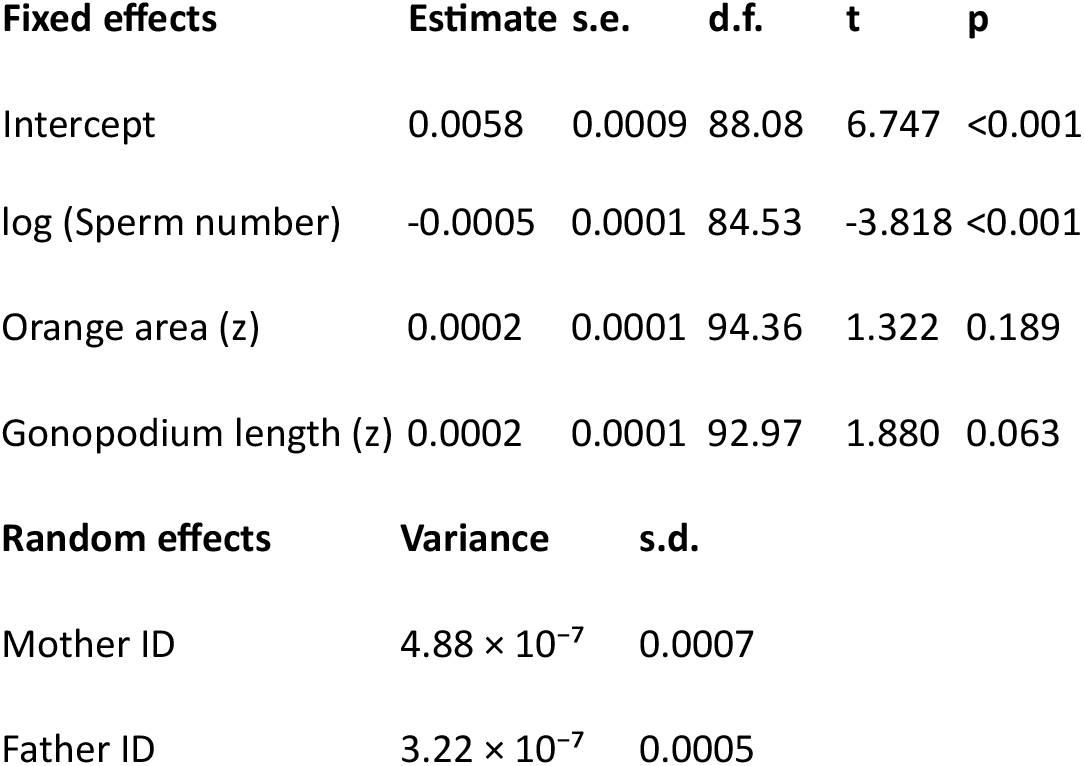
Reduced model excluding sperm velocity. Linear mixed-effects model testing the association between RGR_60_– _120_ and male sexual traits excluding sperm velocity (VAP). Fixed effects include log-transformed sperm number, orange coloration area, and gonopodium length. Maternal and paternal identities were included as random intercepts. Model structure and statistical procedures follow those described in Table S1.

**Table S3.**
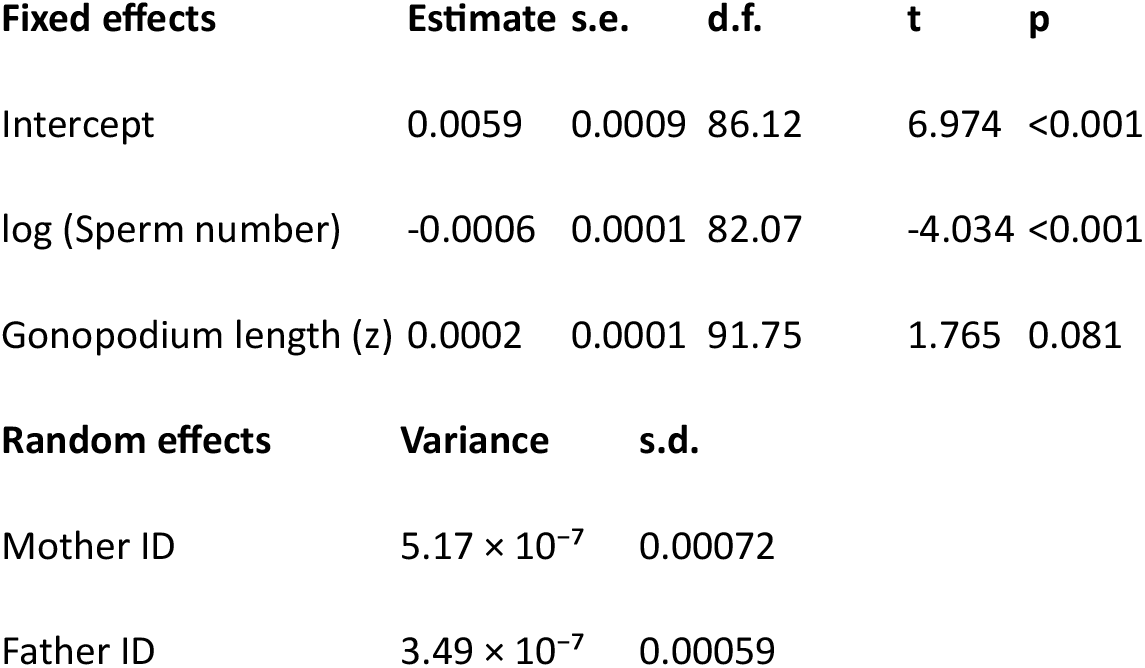
Minimal model including sperm number and gonopodium length. Linear mixed-effects model testing the association between RGR_60_–_120_ and log-transformed sperm number and gonopodium length only. Maternal identity (Mother ID) and paternal identity (Father ID) were included as random intercepts.

### Relationship between social isolation RGR and body size

To assess whether the negative association between RGR_60_–_120_ and sperm production could be explained by differences in adult body size, we examined the relationship between RGR_60_–_120_ and standard length at both 60 and 120 days.

Social isolation growth rate was strongly negatively correlated with body size at 60 days (estimate= -0.00078; *p* < 0.001; Table S4), indicating that males exhibiting higher growth between 60 and 120 days were smaller at the onset of the isolation period. This pattern is consistent with compensatory growth, where initially smaller individuals exhibit accelerated growth rates.

In contrast, RGR_60_–_120_ showed a positive association with body size at 120 days (estimate= 0.00051; *p* < 0.001; Table S5), indicating that individuals with higher growth rates ultimately achieved larger adult body size.

Together, these results indicate that compensatory growth is size-dependent, with smaller males accelerating growth and partially or fully compensating in body size by adulthood. However, despite this compensation, faster growth remains associated with reduced sperm production, supporting the existence of a trade-off between somatic growth and reproductive investment rather than a simple size-mediated effect.

**Table S4.**
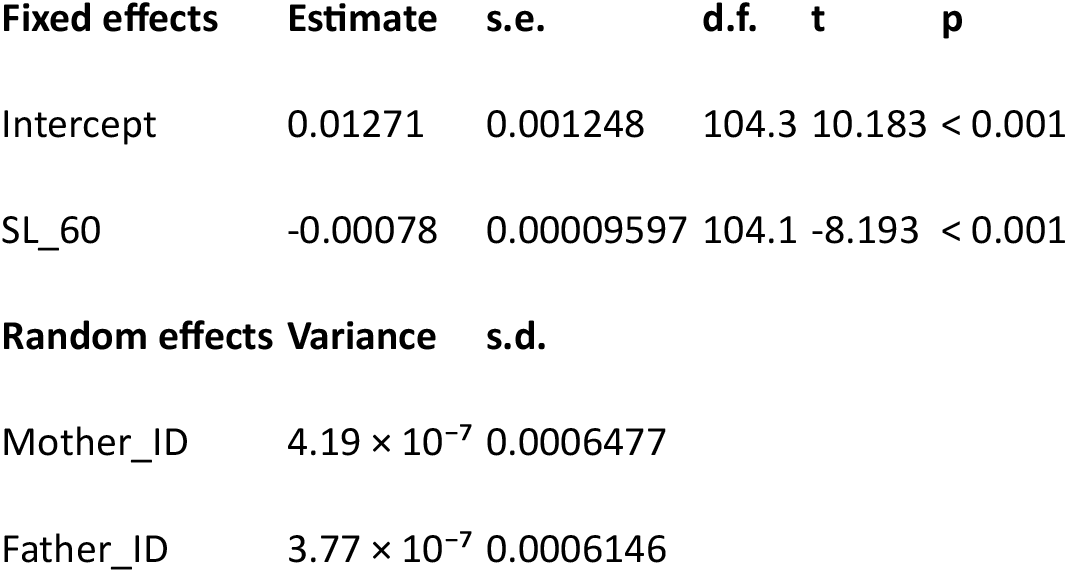
Linear mixed-effects model testing the association between RGR_60_–_120_ and male standard length measured at 60 days of age. Maternal identity (Mother ID) and paternal identity (Father ID) were included as random intercepts. Estimates are reported with standard errors and Satterthwaite-approximated *p*-values.

**Table S5.**
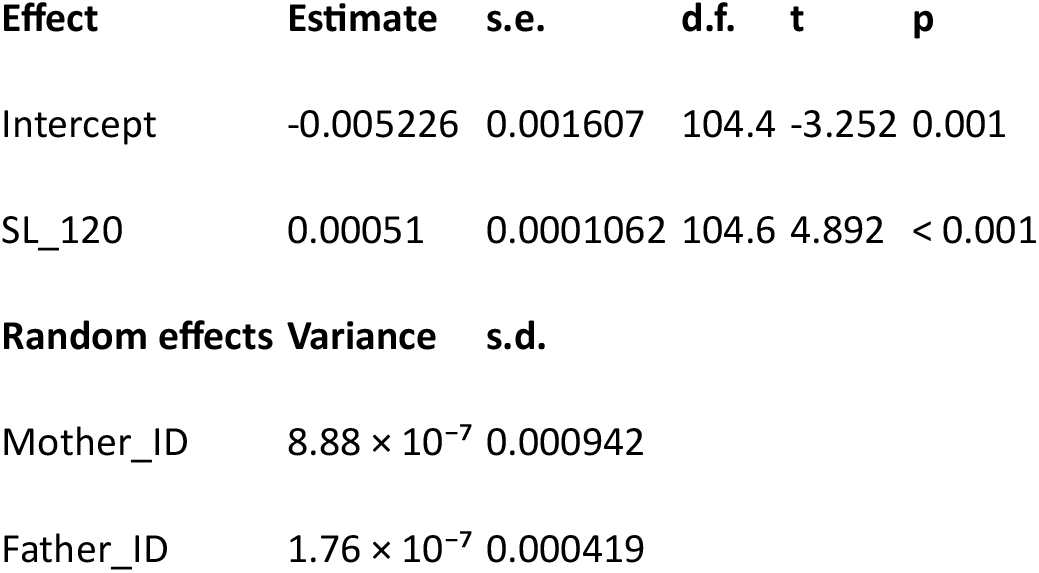
Linear mixed-effects model testing the association between RGR_60_–_120_ and male standard length measured at 120 days of age. Maternal identity (Mother ID) and paternal identity (Father ID) were included as random intercepts. Estimates are reported with standard errors and Satterthwaite-approximated *p*-values.

**Figure S1.**
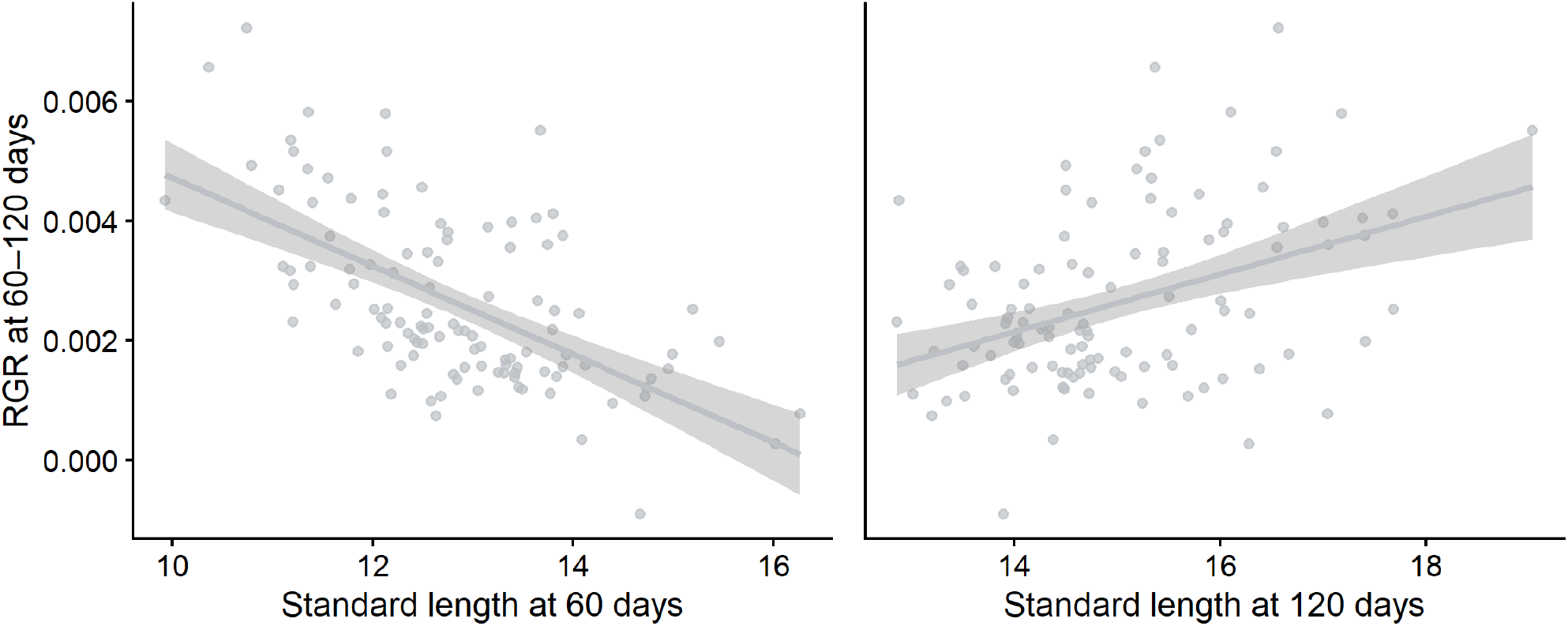
Relationship between RGR_60_–_120_ and body size. Relationship between RGR_60_–_120_ and male body size measured as standard length at 60 days (left panel) and 120 days (right panel). Each point represents an individual male. Solid lines represent fitted linear regressions, and shaded areas indicate 95% confidence intervals.

