## Supplementary Information for "Socially-mediated compensatory growth carries hidden sperm costs in male guppies"

*Affiliations*

### Supplementary Information

#### *Ejaculate assays*

At 120 days of age, males were anesthetized in an MS-222 bath and stripped of sperm immediately after being photographed (see above). Fish were placed under a dissection microscope, the gonopodium swung back and forth and sperm released in a drop of 1% saline solution by applying gentle pressure to the abdomen. Guppy ejaculates are organized in discrete packages (bundles), each containing roughly 21000 sperm cells (Boschetto et al 2011). Ejaculates were photographed with a Canon 450D camera mounted to a light microscope and sperm bundles were counted from pictures using ImageJ. Sperm velocity (µm/s) was measured with a CASA system (Hamilton-Thorne). Three sperm bundles were placed on a multiwell slide coated in 1% polyvinyl acid to prevent adherence, and activated in a medium containing 150 mM KCl and 2 mg/L bovine serum albumin. Velocity was measured as cells swam away from the opening bundle. The sperm tracker produces three measures of velocity (curvilinear, straight line and average path) which are closely corelated in guppies (Gasparini et al 2010). We retained average path velocity (VAP) for analyses.

#### *RGR relationship with pre and postcopulatory traits*

We used multivariate linear mixed-effects models to examine whether $RGR_{60-120}$ was associated with male reproductive traits measured at 120 days of age. Among the examined traits, only sperm number showed a statistically significant association with $RGR_{60-120}$, whereas sperm velocity, coloration, and gonopodium length showed non-significant effects even after removing the least significant predictors in reduced models (Table S1, Table S2, Table S3).

**Table S1.** Linear mixed-effects model testing the association between social isolation RGR ($RGR_{60-120}$) and male sexual traits measured at 120 days of age. Fixed effects include log-transformed sperm number, sperm velocity (VAP), orange coloration area, and gonopodium length. Maternal identity (Mother ID) and paternal identity (Father ID) were included as random intercepts. All continuous predictors were standardized (mean = 0, SD = 1). Estimates are reported with standard errors and Satterthwaite-approximated *p*-values.

**Table S1**

| Fixed effects | Estimate | s.e. | d.f. | t | p |
|---|---|---|---|---|---|
| Intercept | 0.0056 | 0.0009 | 89.13 | 6.509 | <0.001 |
| log (Sperm number) | -0.0005 | 0.0001 | 85.95 | -3.587 | <0.001 |
| VAP (z) | -0.0001 | 0.0001 | 88.36 | -1.247 | 0.216 |
| Orange area (z) | 0.0002 | 0.0001 | 96.07 | 1.652 | 0.102 |
| Gonopodium length (z) | 0.0002 | 0.0001 | 92.23 | 1.786 | 0.077 |
| **Random effects** | **Variance** | **s.d.** | | | |
| Mother ID | $4.68 \times 10^{-7}$ | 0.0006 | | | |
| Father ID | $3.34 \times 10^{-7}$ | 0.0005 | | | |

**Table S2.** Reduced model excluding sperm velocity. Linear mixed-effects model testing the association between $RGR_{60\text{–}120}$ and male sexual traits excluding sperm velocity (VAP). Fixed effects include log-transformed sperm number, orange coloration area, and gonopodium length. Maternal and paternal identities were included as random intercepts. Model structure and statistical procedures follow those described in Table S1.

**Table S2**

| Fixed effects | Estimate | s.e. | d.f. | t | p |
|---|---|---|---|---|---|
| Intercept | 0.0058 | 0.0009 | 88.08 | 6.747 | <0.001 |
| log (Sperm number) | -0.0005 | 0.0001 | 84.53 | -3.818 | <0.001 |
| Orange area (z) | 0.0002 | 0.0001 | 94.36 | 1.322 | 0.189 |
| Gonopodium length (z) | 0.0002 | 0.0001 | 92.97 | 1.880 | 0.063 |
| **Random effects** | **Variance** | **s.d.** | | | |
| Mother ID | $4.88 \times 10^{-7}$ | 0.0007 | | | |
| Father ID | $3.22 \times 10^{-7}$ | 0.0005 | | | |

**Table S3.** Minimal model including sperm number and gonopodium length. Linear mixed-effects model testing the association between $RGR_{60\text{–}120}$ and log-transformed sperm number and gonopodium length only. Maternal identity (Mother ID) and paternal identity (Father ID) were included as random intercepts.

**Table S3**

| Fixed effects | Estimate | s.e. | d.f. | t | p |
|---|---|---|---|---|---|
| Intercept | 0.0059 | 0.0009 | 86.12 | 6.974 | <0.001 |
| log (Sperm number) | -0.0006 | 0.0001 | 82.07 | -4.034 | <0.001 |

**Table S3**

| Fixed effects | Estimate | s.e. | d.f. | t | p |
|---|---|---|---|---|---|
| Gonopodium length (z) | 0.0002 | 0.0001 | 91.75 | 1.765 | 0.081 |

| Random effects | Variance | s.d. |
|---|---|---|
| Mother ID | $5.17 \times 10^{-7}$ | 0.00072 |
| Father ID | $3.49 \times 10^{-7}$ | 0.00059 |

#### *Relationship between social isolation RGR and body size*

To assess whether the negative association between $RGR_{60-120}$ and sperm production could be explained by differences in adult body size, we examined the relationship between $RGR_{60-120}$ and standard length at both 60 and 120 days.

Social isolation growth rate was strongly negatively correlated with body size at 60 days (estimate= -0.00078; $p < 0.001$; Table S4), indicating that males exhibiting higher growth between 60 and 120 days were smaller at the onset of the isolation period. This pattern is consistent with compensatory growth, where initially smaller individuals exhibit accelerated growth rates.

In contrast, $RGR_{60-120}$ showed a positive association with body size at 120 days (estimate= 0.00051; $p < 0.001$; Table S5), indicating that individuals with higher growth rates ultimately achieved larger adult body size.

Together, these results indicate that compensatory growth is size-dependent, with smaller males accelerating growth and partially or fully compensating in body size by adulthood. However, despite this compensation, faster growth remains associated with reduced sperm production, supporting the existence of a trade-off between somatic growth and reproductive investment rather than a simple size-mediated effect.

**Table S4.** Linear mixed-effects model testing the association between $RGR_{60-120}$ and male standard length measured at 60 days of age. Maternal identity (Mother ID) and paternal identity (Father ID) were included as random intercepts. Estimates are reported with standard errors and Satterthwaite-approximated *p*-values.

**Table S4**

| Fixed effects | Estimate | s.e. | d.f. | t | p |
|---|---|---|---|---|---|
| Intercept | 0.01271 | 0.001248 | 104.3 | 10.183 | < 0.001 |
| SL_60 | -0.00078 | 0.00009597 | 104.1 | -8.193 | < 0.001 |

| Random effects | Variance | s.d. |
|---|---|---|
| Mother_ID | $4.19 \times 10^{-7}$ | 0.0006477 |

**Table S4**

| Fixed effects | Estimate | s.e. | d.f. | t | p |
|---|---|---|---|---|---|
| Father_ID | $3.77 \times 10^{-7}$ | 0.0006146 | | | |

**Table S5.** Linear mixed-effects model testing the association between $RGR_{60–120}$ and male standard length measured at 120 days of age. Maternal identity (Mother ID) and paternal identity (Father ID) were included as random intercepts. Estimates are reported with standard errors and Satterthwaite-approximated *p*-values.

**Table S5**

| Effect | Estimate | s.e. | d.f. | t | p |
|---|---|---|---|---|---|
| Intercept | -0.005226 | 0.001607 | 104.4 | -3.252 | 0.001 |
| SL_120 | 0.00051 | 0.0001062 | 104.6 | 4.892 | < 0.001 |
| **Random effects** | **Variance** | **s.d.** | | | |
| Mother_ID | $8.88 \times 10^{-7}$ | 0.000942 | | | |
| Father_ID | $1.76 \times 10^{-7}$ | 0.000419 | | | |

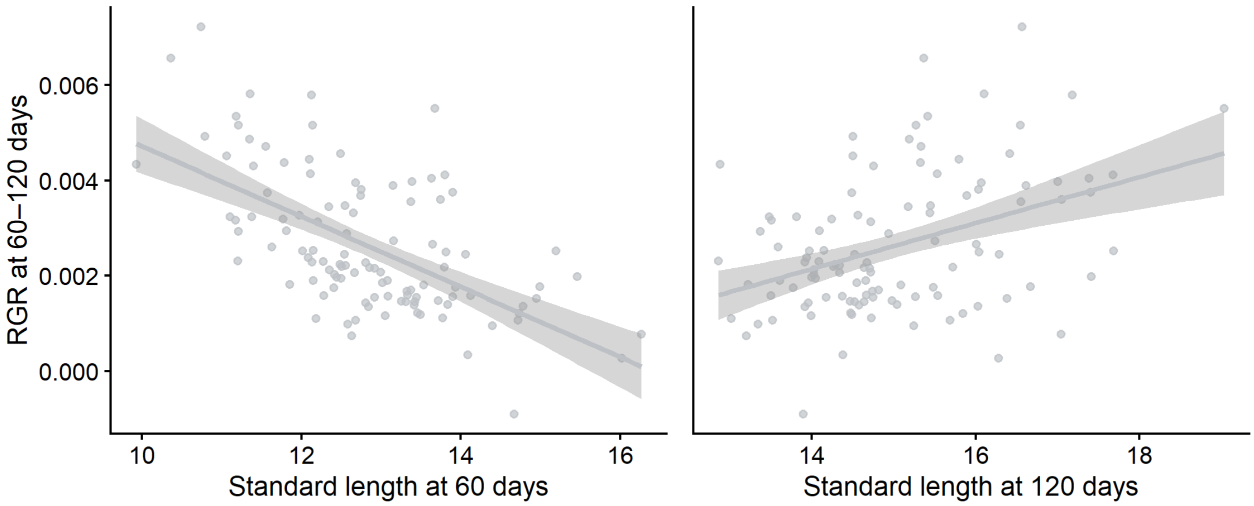


**Figure S1. Relationship between $RGR_{60–120}$ and body size.** Relationship between $RGR_{60–120}$ and male body size measured as standard length at 60 days (left panel) and 120 days (right panel). Each point represents an individual male. Solid lines represent fitted linear regressions, and shaded areas indicate 95% confidence intervals.
